# Bassoon controls synaptic vesicle pools via regulation of presynaptic phosphorylation and cAMP homeostasis

**DOI:** 10.1101/2021.07.22.453360

**Authors:** Carolina Montenegro-Venegas, Debarpan Guhathakurta, Eneko Pina-Fernandez, Maria Andres-Alonso, Florian Plattner, Vesna Lazarevic, Eckart D. Gundelfinger, Anna Fejtova

## Abstract

Neuronal presynaptic terminals contain hundreds of neurotransmitter-filled synaptic vesicles (SVs). The morphologically uniform SVs differ in their release competence segregating into functional pools that differentially contribute to neurotransmission. The presynaptic scaffold bassoon is required for neurotransmission, but the underlying molecular mechanisms are unknown. We report that glutamatergic synapses lacking bassoon featured a decreased SV release competence and increased resting pool of SV as observed by imaging of SV release in cultured neurons. Further analyses in vitro and in vivo revealed a dysregulation of CDK5/calcineurin and cAMP/PKA presynaptic signalling resulting in an aberrant phosphorylation of their downstream effectors synapsin 1 and SNAP25, which are well-known regulators of SV release competence. An acute pharmacological restoration of physiological CDK5 and cAMP/PKA activity fully normalised the SV pools in neurons lacking bassoon. Finally, we demonstrated that CDK5-dependent regulation of PDE4 activity controls SV release competence by interaction with cAMP/PKA signalling. These data reveal that bassoon organises SV pools via regulation of presynaptic phosphorylation and indicate an involvement of PDE4 in the control of neurotransmitter release.

## Introduction

Synapses are contact sites between neurons, where communication between presynaptic and postsynaptic neuron occur by means of electrochemical neurotransmission. Within the presynapse the neurotransmitter is stored in small synaptic vesicles (SVs) and released upon fusion of these SVs with the presynaptic plasma membrane. After their fusion, the membrane and protein components of SVs are retrieved by compensatory endocytosis, SVs are refilled with neurotransmitter and recycled (Sudhof, 2004). This SV cycle ensures long-term structural and functional integrity of the presynaptic compartment (Chanaday et al., 2019; Bonnycastle et al., 2020). SVs, albeit structurally identical, differ in their release competence and kinetics in response to stimuli, resulting in three distinct SV pools (Rizzoli and Betz, 2005; Denker and Rizzoli, 2010; Alabi and Tsien, 2012). The readily releasable pool (RRP) contains vesicles that discharge immediately upon stimulation and correspond morphologically to the docked SVs, which are in contact with the plasma membrane of presynaptic active zone (Schikorski and Stevens, 2001). The recycling pool (RP) is a source of releasable vesicles that can replenish RRP during prolonged stimulation. Both RRP and RP together form the total recycling pool (TRP). A large proportion of SVs is unable to undergo release under physiological conditions and form the reserve pool (ResP) of SVs. The size of RRP and RP at a given synapse decisively shapes its physiological properties. RRP size determines together with release probabilities of individual SVs synaptic release probability, whereas the size of the RP influences the replenishment of SVs during stimulation trains (Alabi and Tsien, 2012). The release competence of SVs and thus their assignment to the functional pools is under the control of intracellular signalling pathways. Cyclin dependent kinase 5 (CDK5)/calcineurin signalling plays a crucial role in the regulation of transition of SVs between the RP and ResP (Kim and Ryan, 2010), while cyclic adenosine monophosphate (cAMP)/cAMP-dependent protein kinase A (PKA) signalling was linked to the control of RRP (Lonart et al., 1998; Nagy et al., 2004). Phosphoproteins of the synapsin (Syn) family are SV-associated proteins that integrate the signalling from kinase and phosphatase of multiple pathways to control plasticity of neurotransmitter release. Syn is critical for the formation and maintenance of the SV cluster above the presynaptic active zone (Pieribone et al., 1995). Depending on its phosphorylation state, Syn oligomerises and interacts with synaptic vesicles and with the actin cytoskeleton, which in turn modulates dynamic SV clustering (Cesca et al., 2010). Syn phosphorylation by cAMP/PKA signalling increases the SV mobilisation. It allows the transition of vesicles from RP to the RRP, but also promotes an increased exchange of SVs between adjacent synaptic varicosities along axons (Menegon et al., 2006; Valente et al., 2012; Patzke et al., 2019; Chenouard et al., 2020). Phosphorylation of Syn by CDK5 enhances its association with actin filaments and mediates a shift of SVs from RP to ResP (Verstegen et al., 2014). Interference with phosphorylation of Syn by PKA and CDK5, respectively, affects the presynaptic short-term and homeostatic plasticity highlighting the importance of dynamic phosphorylation at presynapse for physiological regulation of neurotransmission (Menegon et al., 2006; Valente et al., 2012; Verstegen et al., 2014)

Bassoon (Bsn) is a large scaffolding protein localised exclusively at the active zone of neurotransmitter release (tom Dieck et al., 1998). It plays a key role in the organisation and maintenance of the presynaptic release apparatus (Gundelfinger et al., 2015). Mice with a deletion of the central part of Bsn show generalised seizures and impaired synaptic structure and function (Altrock et al., 2003; Dick et al., 2003). Bsn was shown to coordinate the recruitment of specific presynaptic voltage-gated calcium channels to the active zone via its interaction with RIM-binding proteins (Davydova et al., 2014). Therefore, the Cav2.1 and Cav1.3 channels, respectively, are less abundant in cortico-hippocampal glutamatergic synapses or cochlear and retinal ribbon synapses of Bsn mutants (Frank et al., 2010; Davydova et al., 2014; Babai et al., 2019; Ryl et al., 2021). Investigations of mouse strains expressing shortened or null Bsn alleles detected reduced RRP and slower vesicular replenishment in glutamatergic synapses of cultured hippocampal neurons, cerebellar mossy fibers, photoreceptors, cochlear inner hair cells, and auditory nerve fibers in the absence of functional Bsn (Altrock et al., 2003; Khimich et al., 2005; Frank et al., 2010; Hallermann et al., 2010; Jing et al., 2013; Mendoza Schulz et al., 2014; Babai et al., 2019; Babai et al., 2020; Ryl et al., 2021). The morphological analyses also pointed towards a role of Bsn (and its paralogue Piccolo) in the SV clustering and RRP regulation (Altrock et al., 2003; Mukherjee et al., 2010; Mendoza Schulz et al., 2014; Ackermann et al., 2019; Hoffmann-Conaway et al., 2020). However, the molecular basis for the role of Bsn in the regulation of RRP size and/or release site replenishment remained unclear.

In this study, we investigated the cellular signalling that drives the dynamic regulation of SV recycling using in vivo imaging of hippocampal neurons derived from Bsn mutant mice (Bsn^GT^). We observed a reduced size of RRP and TRP and a significant drop in the overall release competence of SVs in glutamatergic synapses lacking Bsn. These phenotypes were associated with a dysregulation of CDK5/calcineurin balance and cAMP/PKA presynaptic signalling. Acute pharmacological treatment that restored physiological signalling fully normalised the SV release competence in Bsn^GT^ neurons indicating a causal role of aberrant signalling in the loss of release competence in the absence of Bsn. Finally, our data uncovered a key role of CDK5-dependent regulation of PDE4 in the control of SV pools upstream of cAMP/PKA signalling.

## Results

### Deletion of Bsn restricts recycling competence of SVs

Previous studies indicated changes in the size of RRP in the absence of Bsn in various types of synapses. However, to date, the mechanistic understanding of this phenomenon is missing. To monitor SV exocytosis in living neurons, we expressed the pH-sensitive probe, synaptophluorin-tdimer2 (sypHy), in cultured hippocampal neurons that were derived from Bsn^GT^ animals, where Bsn expression was abolished by insertion of a gene trap cassette (Hallermann et al., 2010), and from their WT littermates. In sypHy, pH-sensitive GFP is inserted in the lumenal domain of the integral SV protein synaptophysin. The fluorescence of sypHy is quenched by the acidic milieu of SVs, it increases after SV fusion and exposure to the neutral media and decreases again upon SVs endocytosis and re-acidification, which allows monitoring of SV recycling. The red fluorescent tdimer2 inserted within the cytoplasmic part of synaptophysin permits identification of sypHy expressing cells at rest (Rose et al., 2013). We utilized an established stimulation protocol consisting of 40 action potentials (AP) at 20 Hz to drive fusion of docked SVs (corresponding to RRP) and 900 AP at 20 Hz to release all release-competent SVs (TRP) (Burrone et al., 2006). To prevent reacidification of endocytosed SVs that occurs concomitantly with SV exocytosis during stimulation experiments were performed in the presence of bafilomycin A, a vesicular proton pump inhibitor. Under these conditions, all vesicles that once underwent exocytosis remain visible (Fig 1A, B). We detected significantly lower RRP and TRP in Bsn^GT^ compared with WT neurons (Fig 1A-C, E; RRP: WT 0.13 ± 0.007, Bsn^GT^ 0.08 ± 0.006; TRP: WT 0.41 ± 0.01, Bsn^GT^ 0.29 ± 0.02). Application of ammonium chloride, which alkalizes the SV and un-quenches all sypHy-expressing SVs that do not recycle, revealed a rise in the ResP in Bsn^GT^ (Fig 1A, B; ResP: WT 0.60 ± 0.013, Bsn^GT^ 0.71 ± 0.016). Quantification of the total synaptic intensity of sypHy upon unquenching also confirmed no changes in its synaptic expression levels between Bsn^GT^ and WT neurons (Fig 1A; WT 1061 ± 143.8 AU, Bsn^GT^ 1041 ± 90.21 AU). A clear-cut left shift in the bell-shaped frequency distribution of the response amplitude was evident in Bsn^GT^ upon 40 APs (Fig 1D) and 900APs (Fig 1F) confirming overall lower response magnitude in the Bsn^GT^ neurons compared to WT. This argues against functional inactivation of a fraction of synapses upon deletion of Bsn reported in earlier studies and for overall impairment (Altrock et al., 2003). Collectively, these experiments indicate a role of Bsn in the regulation of release competence of SV and their distribution to RRP and RP.

**Figure 1.**
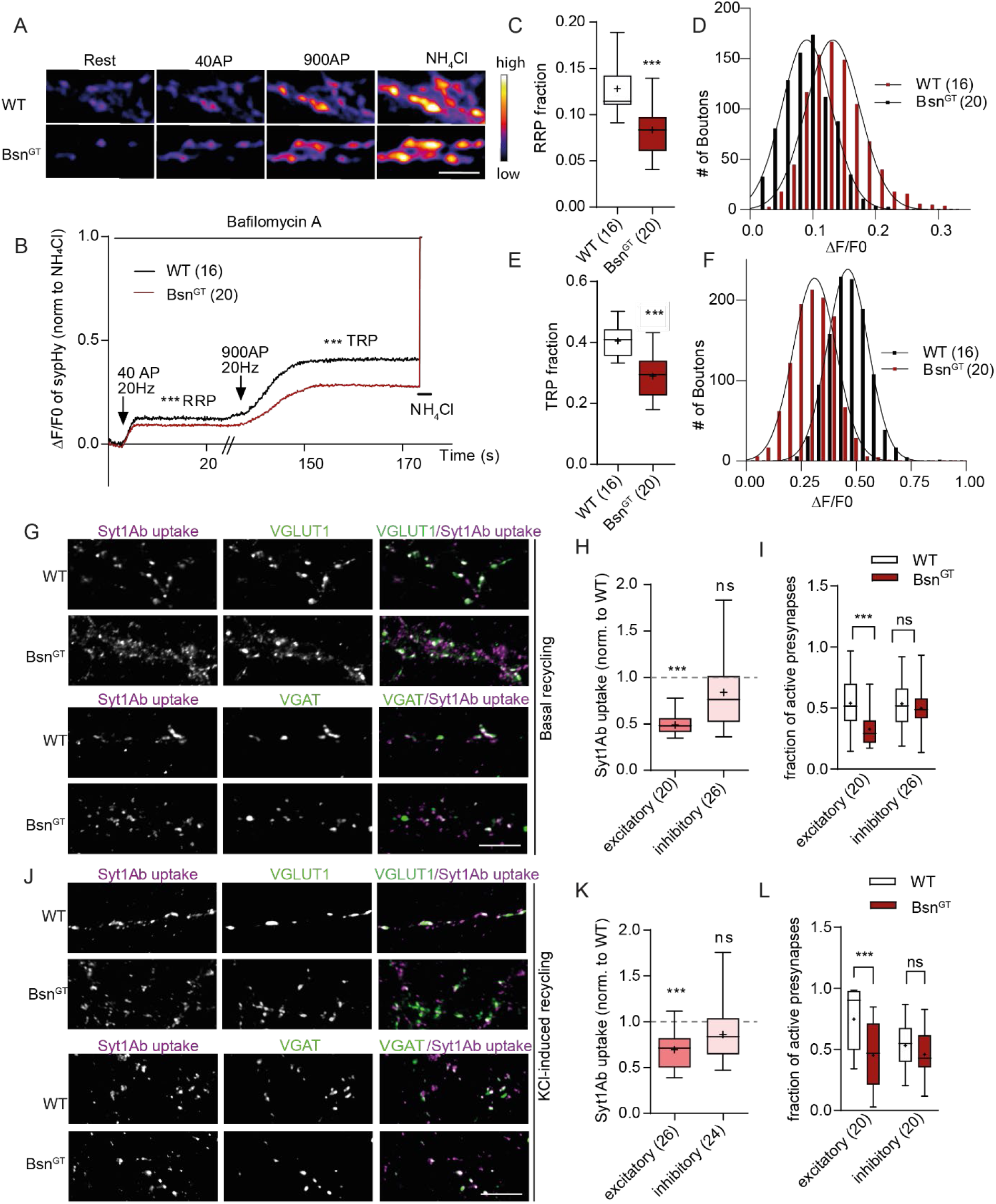
Bassoon deletion affects the release competence of SV at glutamatergic synapse. (A) Representative pseudo colour images of sypHy fluorescence in WT and Bsn^GT^ hippocampal neurons at rest, upon stimulation with 40 and 900 APs at 20Hz (in the presence of bafilomycin A1) and upon application of NH_4_Cl to visualise SVs that were refractory to electrical stimulation. (B) Average traces of the normalised fluorescence change (ΔF/F0) of sypHy in WT (black) and Bsn^GT^ (red) neurons as described in A. Intensities were normalized to the peak of NH_4_Cl response. (C, E) Quantifications of mean RRP and TRP fractions in WT and Bsn^GT^ neurons, respectively. (D, F) Frequency distribution histograms of the response amplitudes of 1050 individual synaptic puncta (from 6 independent experiments) upon 40 AP (D) and 900 AP (F), respectively. The black lines depict superimposed Gaussian fits for each group revealing a shift of the mean value between Bsn^GT^ and WT. (G, J) Representative images of Syt1Ab uptake (magenta) driven by endogenous network activity (G) or after depolarization with 50 mM KCl (J) in hippocampal neurons (18 DIV) from WT and Bsn^GT^ mice. VGLUT1 (green) marks excitatory presynapses (upper panels) and VGAT the inhibitory ones (lower panels). (H-L) Quantification of normalized IF of Syt1Ab uptake (H, K) and of fraction of active (Syt1Ab-labelled) synapses (K, L) performed on experiments illustrated in G and J is shown for each synapse type. In the plots, the interquartile range and median are depicted as boxes, minimal and maximal values as whiskers, and + indicates mean. The sample size n (in parentheses) corresponds to the number of analysed cells. In C and E, the data of 1050 synapses per genotype were processed. Data is obtained from 6 (A-F) or 3 (G-L) independent preparations. Significance was assessed with Student’s t-test *p ≤ 0.05, **p < 0.01, *** p < 0.001. Scale bar is 2 μm in A and 5 μm in G and J.

The hippocampal neuron cultures, used in these experiments contain predominantly excitatory pyramidal cells (Kaech et al., 2012). To assess the SV recycling specifically in excitatory and inhibitory neurons we combined labelling of active synapses in living neurons using a fluorophore-coupled anti-synaptotagmin1 antibody (Syt1Ab) with post-fixation synapse-type specific immunostaining. This Syt1Ab antibody recognizes the luminal domain of the integral SV protein Syt1. If added to the media of living cells, it can access its epitope only after fusion of SV with the plasma membrane, when the extracellular media contacts the lumen of SV (Kraszewski et al., 1995; Lazarevic et al., 2011; Lazarevic et al., 2017). First, we performed Syt1Ab labelling in living neurons without any further manipulations to monitor SV recycling driven by endogenous neuronal activity (Fig 1G). The excitatory synapses were assessed as Syt1Ab-labeled puncta positive for synapsin and vesicular glutamate transporter 1 (VGLUT1) immunoreactivity, and inhibitory synapses as puncta co-labelled for synapsin and vesicular GABA transporter (VGAT) (Fig 1G). Syt1Ab uptake driven by basal network activity was significantly lower in Bsn^GT^ excitatory presynapses, while mild, but not significant decrease was seen in inhibitory buttons (Fig 1G, H; exc: WT 1.00 ± 0.08, Bsn^GT^ 0.49 ± 0.02; inh: WT 1.00 ± 0.07, Bsn^GT^ 0.84 ± 0.08). To assess the total recycling pool in both synapse types, we performed Syt1-Ab labelling during depolarization induced by brief application of 50 mM KCl, which leads to the release of the total recycling pool of SV (TRP) (Harata et al., 2001). Depolarization-induced Syt1Ab uptake was significantly decreased in the excitatory, but not in the inhibitory synapses in neurons derived from Bsn^GT^ animals (Fig 1J, K; exc: WT 1.00 ± 0.07, Bsn^GT^ 0.69 ± 0.04; inh: WT 1.00 ± 0.06, Bsn^GT^ 0.86 ± 0.07).

Post fixation labelling of neurons with a general synaptic marker after Syt1Ab uptake also enabled us to quantify the number of active vs. silent presynapses, i.e. presynapses competent of SV release and recycling under basal network activity or upon chemical depolarization (Moulder et al., 2010). The fraction of active glutamatergic synapses, defined as VGLUT1-positive puncta with detectable immunofluorescence (IF) of Syt1Ab uptake, was reduced by 40% in Bsn^GT^ neurons under basal conditions, indicating a higher proportion of silent (i.e. release incompetent) presynapses (Fig 1 I; WT 0.54 ± 0.05, Bsn^GT^ 0.33 ± 0.03). The number of glutamatergic synapses capable of release upon KCl-induced depolarization was also decreased in Bsn^GT^ neurons compared to the WT (Fig 1L; WT 0.75 ± 0.05, Bsn^GT^ 0.45 ± 0.05). No significant differences in number of release-competent inhibitory, i.e. VGAT and synapsin-positive, synapses were detected (Fig 1I, L; basal: WT 0.55 ± 0.04, Bsn^GT^ 0.50 ± 0.04; KCl: WT 0.53 ± 0.03, Bsn^GT^ 0.46 ± 0.04). These analyses indicate that loss of Bsn affects the SV release competence predominantly at glutamatergic synapses.

### CDK5 activity is increased in Bsn^GT^ and uncoupled from regulation by neuronal activity

CDK5 plays an important role in the regulation of release competence of SV, especially via dynamic recruitment of SVs from the RP and to the ResP (Kim and Ryan, 2010, 2013; Verstegen et al., 2014). To assess the potential involvement of CDK5 in the shift of SV from recycling to resting pool that we observed in Bsn^GT^ neurons, we pharmacologically interfered with the activity of this enzyme in WT and Bsn^GT^ neurons. In accordance with the reported role of CDK5, we observed a significant increase in the fraction of RRP and TRP in WT neurons after acute inhibition of CDK5 with roscovitine (100 μM, 30 min). The analyses of data revealed that roscovitine increased RRP by 68% and TRP by 57% in Bsn^GT^, but only by 38% (RRP) and 29% (TRP) in WT (Fig 2A, B, E; WT vs. Bsn^GT^; RRP 138 ± 9 vs. 168 ± 10%; TRP: 129 ± 5 vs. 157 ± 7%). Importantly, the roscovitine treatment restored RRP and TRP fractions in Bsn^GT^ neurons to the levels similar to treated WT neurons (Fig 2B, C, D; RRP: WT 0.13 ± 0.01, WT + roscovitine 0.18 ± 0.01, Bsn^GT^ 0.09 ± 0.01, Bsn^GT^ + roscovitine 0.15 ± 0.01; TRP: WT 0.40 ± 0.01, WT + roscovitine 0.51 ± 0.02, Bsn^GT^ 0.29 ± 0.02, Bsn^GT^ + roscovitine 0.45 ± 0.02). This indicates that an elevated activity of CDK5 possibly contributes to the changes in SV pools in Bsn^GT^ neurons.

**Figure 2.**
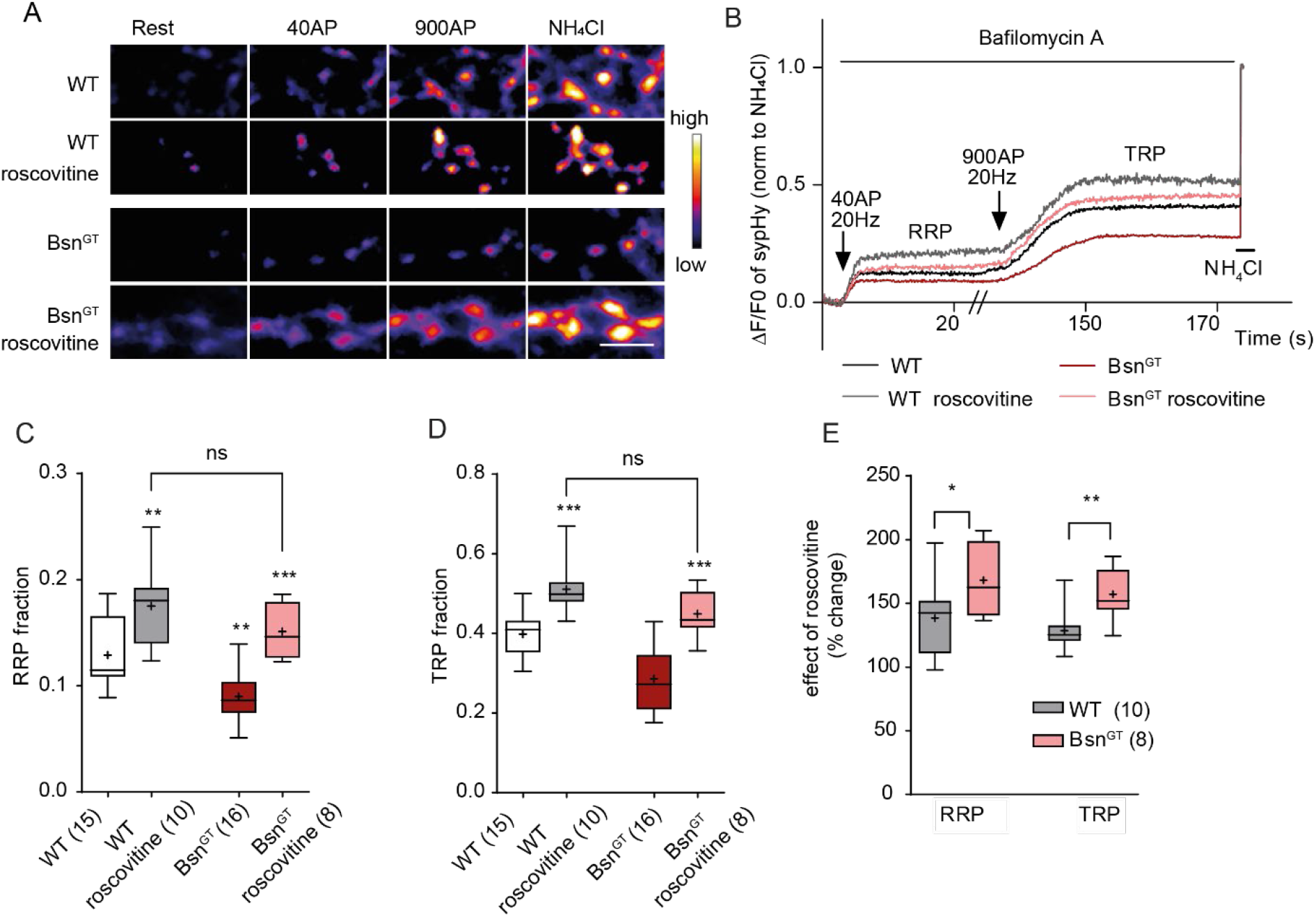
Inhibition of CDK5 activity normalises SV pools in Bsn^GT^ neurons. Representative pseudo colour images (A) and average traces (B) of sypHy fluorescence plotted for WT and Bsn^GT^ neurons treated with a CDK5 inhibitor roscovitine (100μM, gray and pink trace) or vehicle (black and red trace) for 30 minutes before stimulation with 40 and 900 APs in the presence of bafilomycin A. (C, D) Plots show mean values of RRP (C) and TRP (D) fraction for both genotypes before and after treatment. (E) Roscovitine treatment has a significantly higher effect on RRP and TRP in Bsn^GT^ neurons compared to WT. The number of imaging experiments done on 4 independent cell preparations is given in brackets. In the plots, the interquartile range and median are depicted as boxes, minimal and maximal values as whiskers, and + indicates mean. Significance was assessed by one-way ANOVA with Tukey’s post hoc test (C, D) and by Student’s t-test (E) *p≤0.05, **p < 0.01, ***p< 0.001. Scale bar is 2μm.

An important substrate of CDK5 in the context of regulation of SV recycling is the Ser551 of SV-associated protein synapsin1 (Verstegen et al., 2014). Phosphorylation at Ser551 of Syn1 (pSer551Syn1) enhances the binding of Syn1 to F-actin, promotes a shift of SVs to the ResP, and thereby, it restricts SV recycling (Fig 3C) (Verstegen et al., 2014). To address whether deregulation of CDK5-dependent pSer551Syn1 underlies the changes in SV pools in Bsn^GT^ neurons, we utilized the previously published phospho-specific antibody (Verstegen et al., 2014). We detected significantly higher levels of pSer551Syn1 in P2 fractions of hippocampal lysates from Bsn^GT^ mice than in WT, while the total expression levels of Syn1 were unchanged (Fig 3A, B; WT 1.00 ± 0.06, Bsn^GT^ 1.59 ± 0.14). Next, we used this antibody to visualize abundance of pSer551Syn1 in individual synapses in cultured WT and Bsn^GT^ neurons. We detected a significantly higher pSer551Syn1 labelling in synapses of neurons from Bsn^GT^ compared to the WT, while the total expression of Syn did not differ between genotypes (Fig 3D, E; WT 1.00 ± 0.06, Bsn^GT^ 1.55 ± 0.08). Importantly, the pSer551Syn1 immunoreactivity was dramatically decreased in WT terminals upon treatment with roscovitine, confirming the specificity of the antibody (Fig 3D, E; 0.43 ± 0.03). This result indicates an increased CDK5-mediated phosphorylation of Ser551 of Syn1 in Bsn^GT^ mice.

**Figure 3.**
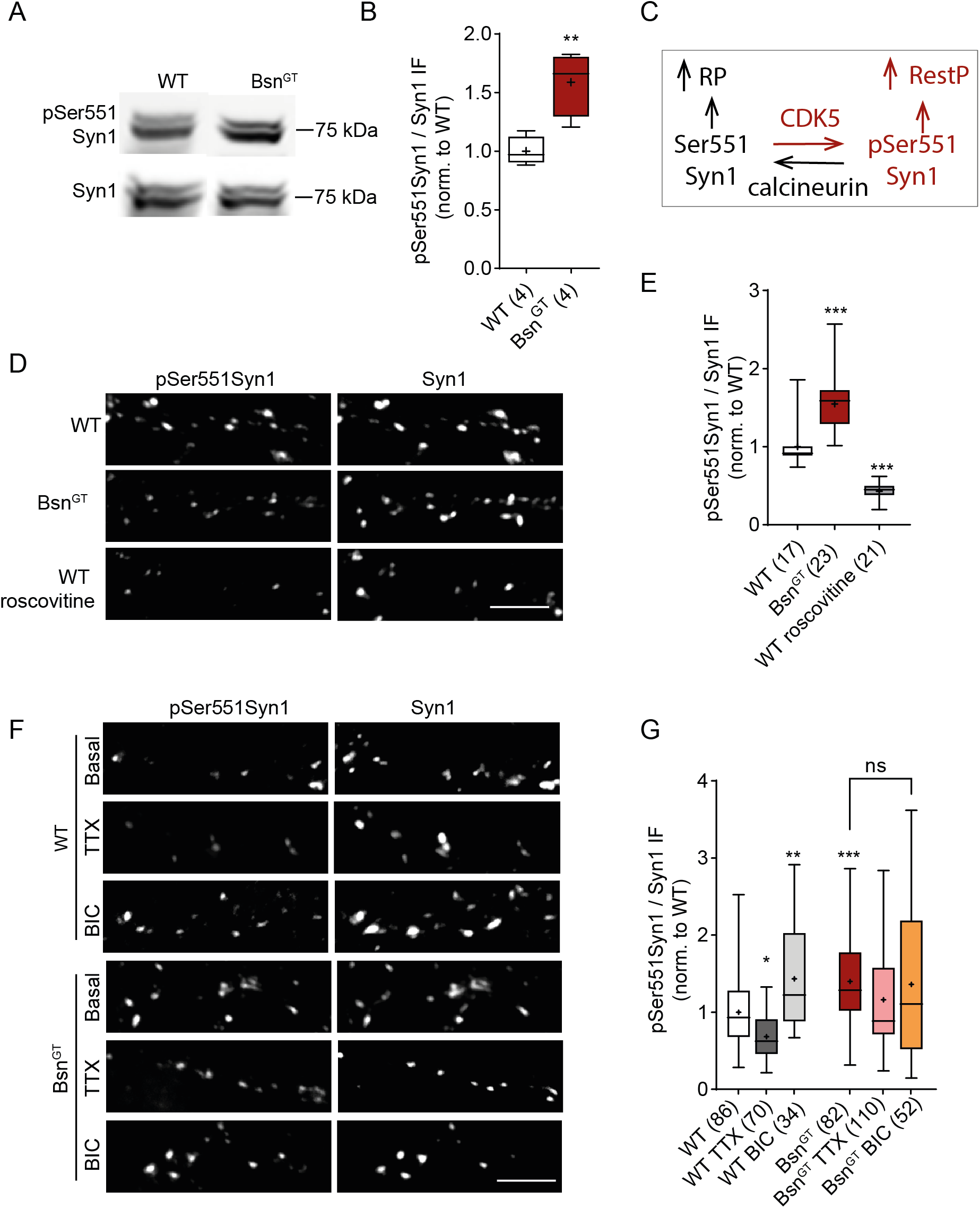
CDK5-dependent phosphorylation of Ser551 of Syn1 is increased in Bsn^GT^ in vivo and in vitro and not regulated by neuronal activity in vitro. A) Representative immunoblot with antibody against pSer551 of Syn1 (pSer551Syn1) by CDK5 and against total Syn1 on P2 fractions prepared from hippocampal tissue lysates of WT and Bsn^GT^ mice. B) Quantification of pSer551Syn1 / Syn1 ratio in Bsn^GT^ and WT (C) Scheme depicts the regulation of phosphorylation of Ser551 of Syn1 by CDK5/calcineurin balance that mediates recruitment of SVs from TRP to the ResP. Changes in this signalling detected in Bsn^GT^ neurons are depicted in red. (D) Representative images of 19 DIV cultured hippocampal neurons from WT and Bsn^GT^ immunostained for pSer551Syn1 and total Syn1. (E) Quantification of fluorescence intensity of pSer551Syn1 in synapses from WT, Bsn^GT^ and WT cells treated with roscovitine. (F) Representative images of hippocampal WT and Bsn^GT^ neurons at baseline activity, upon network activity silencing (TTX, 1 μM, 72 h) or in conditions of increase activity (BIC, 30 μM, 48 h) G) Quantification of experiment in F. In the plots the interquartile range and median are depicted as boxes, minimal and maximal values as whiskers and + indicates mean. The sample size n (in brackets) corresponds to the number of independently processed animals in B or to the number of independent visual fields (D, E) quantified from two (D) and four (E) independent cultures per condition. For statistics, student’s t-test in B and one-way ANOVA with Tukey’s post hoc test in E and G were used, *p≤0.05, **p < 0.01, ***p< 0.001. Scale bar is 5 μm in D and F.

The kinase activity of CDK5 at the presynapse is under the control of neuronal activity (Kim and Ryan, 2010). Bsn deletion leads to epileptic seizures (Altrock et al., 2003) and it is therefore possible that increased endogenous network activity triggers higher activation of CDK5 in Bsn^GT^. To address this, we treated both WT and Bsn^GT^ neurons with tetrodotoxin (TTX) and bicuculline (BIC), which decreases or increases, respectively, phosphorylation at Ser551 of Syn1 in cultured neurons (Verstegen et al., 2014). In line with published data, TTX treatment (1 μM, 72 h) significantly decreased, while application of BIC (30μM, 48 hrs) significantly elevated Ser551Syn1 phosphorylation in WT neurons Fig 3F, G; WT 1.00 ± 0.05, WT + TTX 0.68 ± 0.03, WT + BIC 1.43 ± 0.10). However, neither of treatments affected the pSer551Syn1 in Bsn^GT^ neurons (Fig 3F, G; Bsn^GT^ 1.40 ± 0.07, Bsn^GT^ + TTX 1.16 ± 0.06, Bsn^GT^ +BIC 1.36 ± 0.13) These data indicate that increased activity of CDK5 at presynapse contributes to the changes in SV pools in Bsn^GT^ neurons. Moreover, they revealed a failure in neuronal activity-dependent regulation of presynaptic CDK5 activity in the absence of Bsn.

### Regulation of RRP by PKA and calcineurin is dysregulated upon Bsn deletion

Calcineurin is the Ser/Thr phosphatase that counteracts the action of CDK5 in the regulation of neurotransmitter release (Kim and Ryan, 2010, 2013). To address the role of calcineurin in the dysregulation of SVs pools in Bsn^GT^ neurons, we applied the calcineurin inhibitor FK506 (1 μM) to neurons 0.5 h prior to imaging of SV pools. This treatment significantly decreased TRP in WT, but had no additional effect on Bsn^GT^ neurons, resulting in the same residual TRP in both genotypes (Fig 4A, B, D; TRP: WT 0.41± 0.01, WT+ FK506 0.27 ± 0.02, Bsn^GT^ 0.29 ± 0.02, Bsn^GT^+ FK506 0.27 ± 0.02). These data are in line with the role of the CDK5/calcineurin balance in the regulation of TRP in WT and with a shift in this balance towards increased CDK5 activity in Bsn^GT^ neurons (Fig 3C). Interestingly, calcineurin inhibition tended to mildly decrease the RRP size in WT, but it significantly increased RRP size in Bsn^GT^ neurons and in fact normalized the impairment in RRP size seen in the absence of Bsn (Fig 4A-C; WT 0.11 ± 0.00, WT + FK506 0.09 ± 0.01, Bsn^GT^ 0.08 ± 0.01, Bsn^GT^ + FK506 0.11 ± 0.01). This observation suggests that depending on the presence of Bsn calcineurin differentially regulates the RRP.

**Figure 4.**
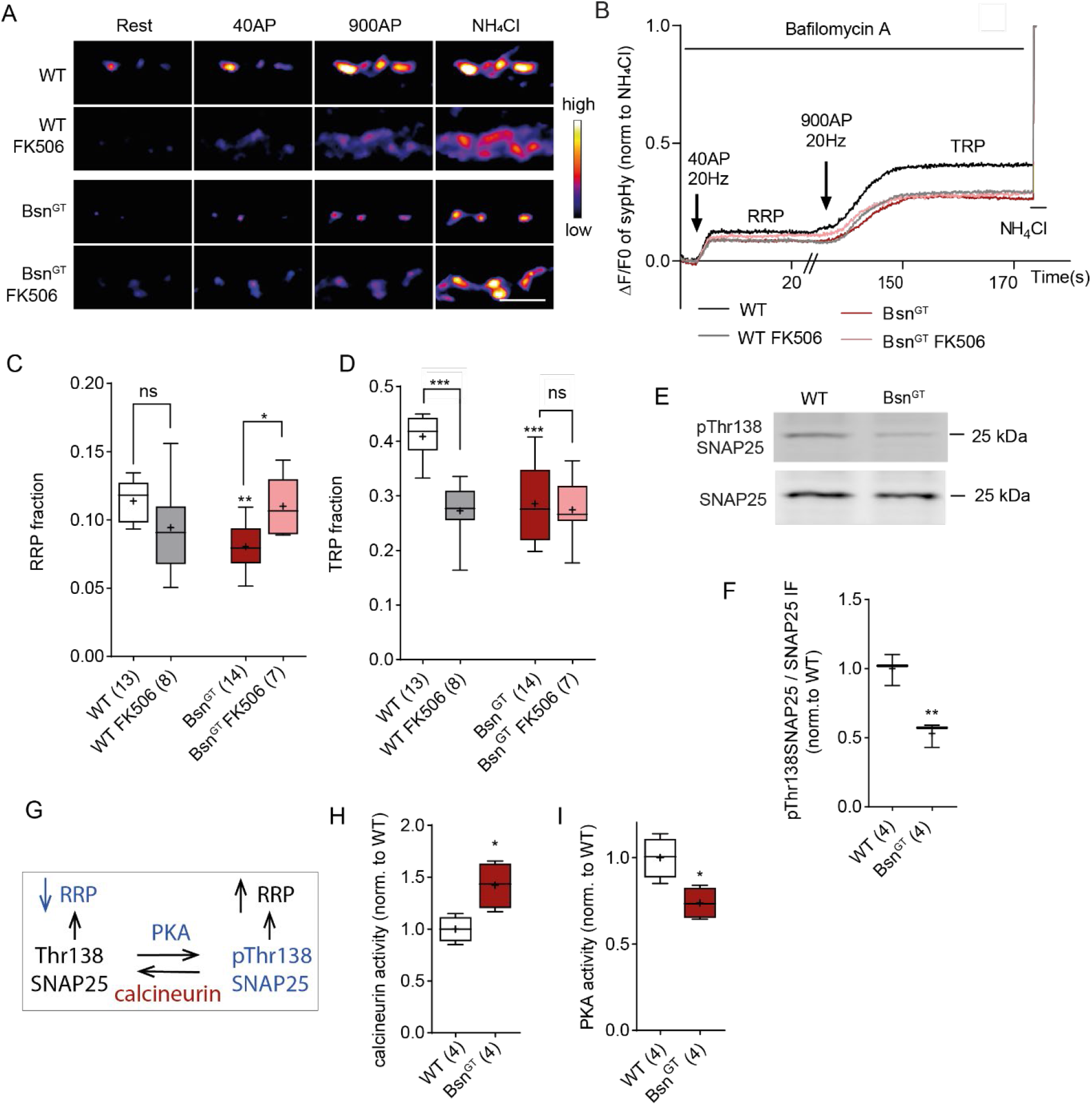
Decreased PKA and increased calcineurin activity in Bsn^GT^ neurons. Representative pseudo colour images (A) and average traces (B) of sypHy fluorescence plotted from WT and Bsn^GT^ hippocampal neurons treated with calcineurin inhibitor FK506. (C, D) Quantification of the RRP and TRP fractions in WT and Bsn^GT^ neurons from experiment in A. (E) Representative Western blot on P2 fractions prepared from hippocampal tissue of WT and Bsn^GT^ mice probed with antibodies against SNAP25 phosphorylated onThr138 and against total SNAP25. F) Quantification of blot in F shows lower pThr138SNAP25 to total SNAP25 IF ratio in Bsn^GT^ compared to WT. (G) Scheme illustrates the PKA/calcineurin-dependent regulation of SNAP25 phosphorylation on its Thr138 that promotes enlargement of RRP via an increase in SV release competence. The changes detected in Bsn^GT^ are depicted in red for elevated and in blue for lowered abundance/activity. (H, I) Direct activity measurements reveal significant increase in calcineurin and drop in PKA activity in samples prepared from hippocampal tissue of Bsn^GT^ mice as compared to their WT littermates. In all plots, the interquartile range and median are depicted as boxes, minimal and maximal values as whiskers, and + indicates mean. The sample size is given as the number in brackets for each quantification and reflect the number of analysed imaging experiments done on 3 independently prepared cultures per genotype in B-D and number of animals per genotype used for WB or enzyme activity measurements in F, H and I. Statistical significance was assessed using one-way ANOVA with Tukey’s post hoc test for C and D and student’s t-test in F, H and I; significance is depicted as follows: *p≤0.05, **p < 0.01, ***p< 0.001. Scale bar is 2 μm.

PKA is a critical regulator of RRP in chromaffin cells with Thr138 of SNAP25 (pThr138SNAP25) being the known target of its action in this context (Fig 4G) (Risinger and Bennett, 1999; Nagy et al., 2004). Phosphatase activity of calcineurin antagonizes this effect of PKA by dephosphorylation of pThr138SNAP25 (Nagy et al., 2004). Thus, we compared the phosphorylation of SNAP25 in WT and Bsn^GT^. Using quantitative immunoblots, we detected significantly lower immunoreactivity for pThr138 SNAP25 in the membrane fraction P2 prepared from Bsn^GT^ hippocampal lysates, while the total expression of SNAP25 was unchanged between genotypes (Fig 4E, F; WT 1.00 ± 0.07, Bsn^GT^ 0.53 ± 0.05). Subsequent direct measurement of enzyme activity showed significantly increased calcineurin phosphatase activity and decreased PKA activity in the hippocampal lysates from Bsn^GT^ (Fig 4H, I; calcineurin: WT 1.00 ± 0.06, Bsn^GT^ 1.42 ± 0.11; PKA: WT 1.00 ± 0.06, Bsn^GT^ 0.73 ± 0.05). Thus, these results indicate that lack of Bsn induces an aberrant presynaptic activity of calcineurin and PKA.

### PKA-dependent regulation of SV pools is shifted and insensitive to forskolin in Bsn^GT^

To test the contribution of PKA in the decreased RRP and TRP size, we compared the effect of pharmacological manipulation of PKA activity on SV recycling visualized by imaging of sypHy fluorescence in WT and Bsn^GT^ neurons. First, we performed measurements in cells of both genotypes treated with H89 (10 μM, 1 hr), a potent inhibitor of PKA. The analysis revealed a significant drop in both RRP and TRP in WT neurons (Fig 5A-D; RRP: WT 0.13 ± 0.01, WT + H89 0.08 ± 0.01; and TRP: WT 0.40 ± 0.01, WT + H89 0.29 ± 0.03). This is in line with a requirement for PKA activity in the regulation of RRP and TRP. The same treatment had a minor effect on RRP and TRP in Bsn^GT^, indicating very low PKA activity in the absence of Bsn (Fig 5A-D; RRP: Bsn^GT^ 0.09 ± 0.01, Bsn^GT^ + H89 0.07 ± 0.01, and TRP: Bsn^GT^ 0.29 ± 0.02, Bsn^GT^ + H89 0.27 ± 0.01). Next, we applied forskolin (50 μM, 1h), a potent nonselective activator of adenylyl cyclases (ACs). ACs are the enzymes catalysing conversion of ATP to cAMP and are therefore upstream of PKA activation. In line with the central role of PKA in the regulation of RRP and TRP, forskolin application induced a significant increase in RRP and TRP in WT neurons. Strikingly, forskolin application had no effect on RRP and TRP in the Bsn^GT^ neurons, indicating a defect in forskolin-induced AC activation in the absence of Bsn (Fig 5 A-D; RRP: WT+ forskolin 0.17 ± 0.01; Bsn^GT^ + forskolin 0.10 ± 0.01; TRP: WT + forskolin 0.50 ± 0.01, Bsn^GT^+ forskolin 0.32 ± 0.03).

**Figure 5.**
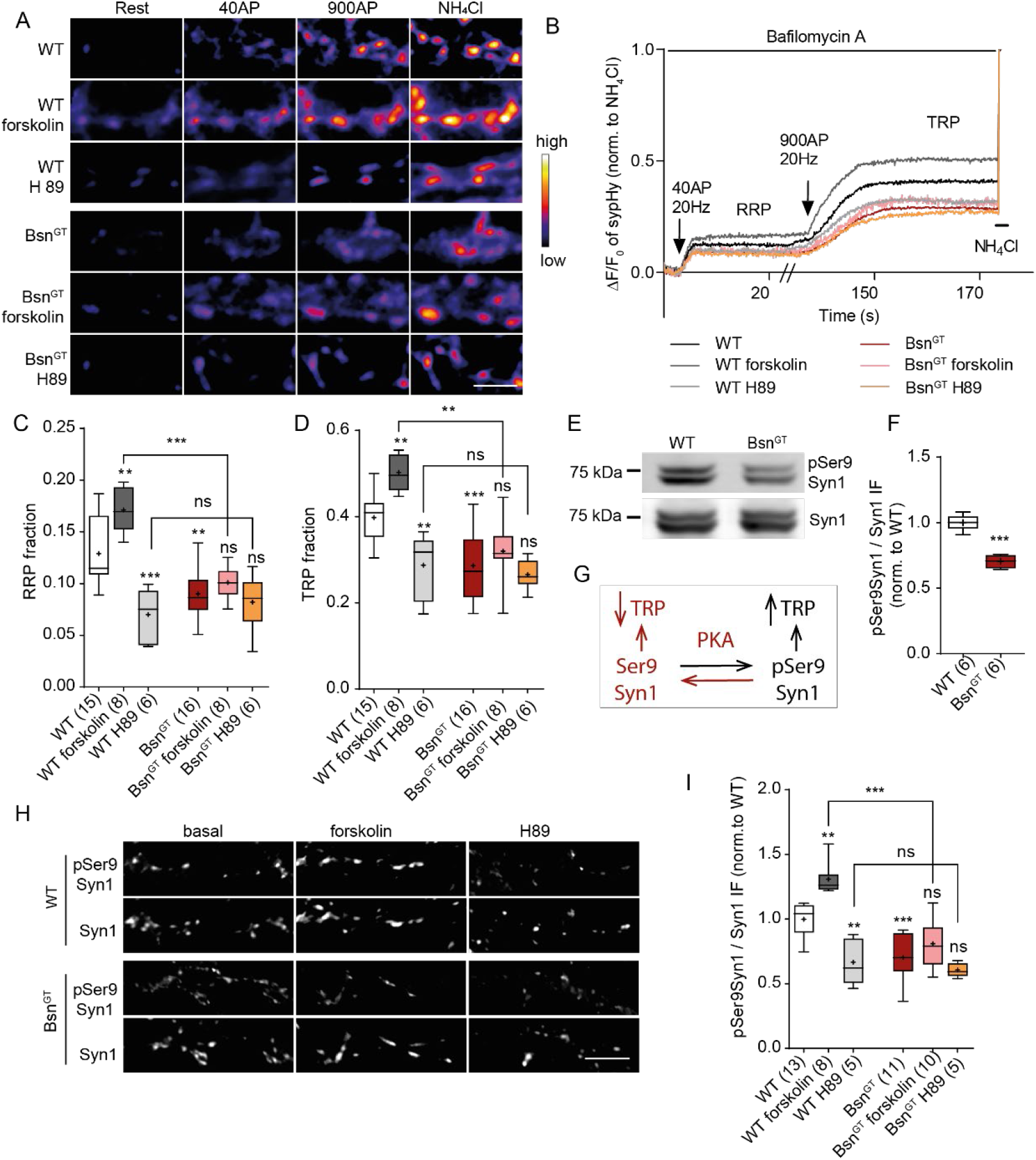
PKA-dependent phosphorylation of Syn1 is decreased and insensitive to forskolin in Bsn^GT^. (A) Representative pseudo colour images (A) and average traces (B) of sypHy fluorescence plotted for WT and Bsn^GT^ neurons without treatment (black and red) and upon treatment with adenylyl cyclase activator forskolin (gray and pink) or PKA inhibitor H89 (light grey and orange). (C, D) Quantification of the RRP and TRP fraction from experiment in A. (E) Representative immunoblot with antibody against pSer9Syn1 and total Syn1 on hippocampal tissue from WT and Bsn^GT^ mice (F) Quantification of blots from E. (G) Scheme illustrates the PKA-dependent phosphorylation of Syn1 on Ser9 that promotes recruitment of SVs to the recycling pool. Changes in this signalling confirmed in Bsn^GT^ are depicted in red. (H) Representative images of hippocampal neurons from WT and Bsn^GT^ mice without treatment and treated with forskolin or H89 labelled with antibodies against pSer9Syn1 and total Syn1. (I) Quantification of staining in H. In the plots, the interquartile range and median are depicted as boxes, minimal and maximal values as whiskers and + indicates mean. The sample size is given in brackets and corresponds to the number of analysed independent imaging experiments performed on three independently prepared culture batches in (C and D), samples prepared from individual animals in (E) or quantified independent visual fields obtained from 2 independent culture preparations in (I). The statistical significance was assessed in C, D and I using one-way ANOVA with Tukey’s post hoc test and in F using Student’s t-test as is depicted in graphs as *p≤0.05, **p < 0.01, ***p< 0.001. Scale bar is 2 μm in A and 5 μm in H.

The regulation of RRP and TRP by PKA relies largely on PKA-dependent phosphorylation of Ser9 of Syn1(pSer9Syn1), which regulates the association of SV with actin filaments and their recruitment to the release sites (Fig 5G) (Menegon et al., 2006; Valente et al., 2012). In line with the decreased presynaptic PKA activity in Bsn^GT^ neurons the phosphorylation of pSer9Syn1 was significantly decreased in the P2 membrane fraction of hippocampal lysates from Bsn^GT^ mice compared to WT (Fig 5E, F; WT 1.00 ± 0.02, Bsn^GT^ 0.70 ± 0.02). Analyses of immunoreactivity for pSer9Syn1 in individual synapses further confirmed that phosphorylation of this critical residue correlates with the functional impairment of RRP and TRP in Bsn^GT^ neurons. Specifically, the pSer9Syn1 immunoreactivity was significantly lower in Bsn^GT^ neurons under basal conditions (Fig 5H, I; WT 1.00 ± 0.04, Bsn^GT^ 0.70± 0.05) and the treatment with H89 decreased pSer9Syn1 immunoreactivity in WT neurons to the levels measured in Bsn^GT^ (Fig 5H, I; WT + H89 0.67 ± 0.08, Bsn^GT^ + H89 0.61 ±0.02). Application of forskolin resulted in increased pSer9Syn1 immunoreactivity in WT neurons, but had no significant effect in Bsn^GT^ (Fig 5I, J; WT+ forskolin 1.31 ± 0.04, Bsn^GT^ + forskolin 0.81 ± 0.06). Collectively, these data indicate that dysregulation of PKA underlies the decrease in RRP and TRP in Bsn^GT^ neurons. Moreover, since the application of AC activator forskolin cannot normalize the dysregulation of SV pools in Bsn^GT^, they indicate an uncoupling of AC-cAMP-PKA signalling in the absence of Bsn.

### PDE4 controls SV pools and is dysregulated in Bsn^GT^

The failure of forskolin to potentiate SV recycling in Bsn^GT^ might be due to the loss of PKA responsiveness to cAMP or to an aberrant cAMP metabolism. To address this, we utilised an analogue of cAMP, Sp-isomer of N6-Benzoyladenosine-3’,5’-cyclic monophosphorothioate (6-Bnz-cAMPS) that acts as a specific activator of PKA and that is resistant to hydrolysis by phosphodiesterases (PDEs). We treated cultured WT and Bsn^GT^ neurons with 6-Bnz-cAMPS (50 μM, 1 h) and analysed the effect of this treatment on SV recycling (Fig 6A, B). The treatment increased RRP by 42% and TRP by 31% in WT, but had a significantly larger effect in Bsn^GT^ neurons, increasing RRP by 78% and TRP by 56% (Fig 6E). Importantly, application of 6-Bnz-cAMPS completely normalized the differences in SV pools measured in untreated Bsn^GT^ neurons (Fig 6A-D; RRP: WT 0.12 ± 0.006, WT + Sp-6-Bnz-cAMPS 0.17 ± 0.01, Bsn^GT^ 0.08 ± 0.01, Bsn^GT^ + Sp-6-Bnz-cAMPS 0.15 ± 0.01, and TRP: WT 0.38 ± 0.02, WT + Sp-6-Bnz-cAMPS 0.50 ± 0.03, Bsn^GT^ 0.29 ± 0.01, Bsn^GT^ + Sp-6-Bnz-cAMPS 0.45 ± 0.03). Thus, the decreased release and recycling competence of SV in Bsn^GT^ is likely due to the lower cAMP abundance in the absence of Bsn. Indeed, a direct measurement revealed lower cAMP concentration in the hippocampal tissue of Bsn^GT^ mice compared to their WT littermates (Fig 6F; WT 1.00 ± 0.09, Bsn^GT^ 0.78 ± 0.03).

**Figure 6.**
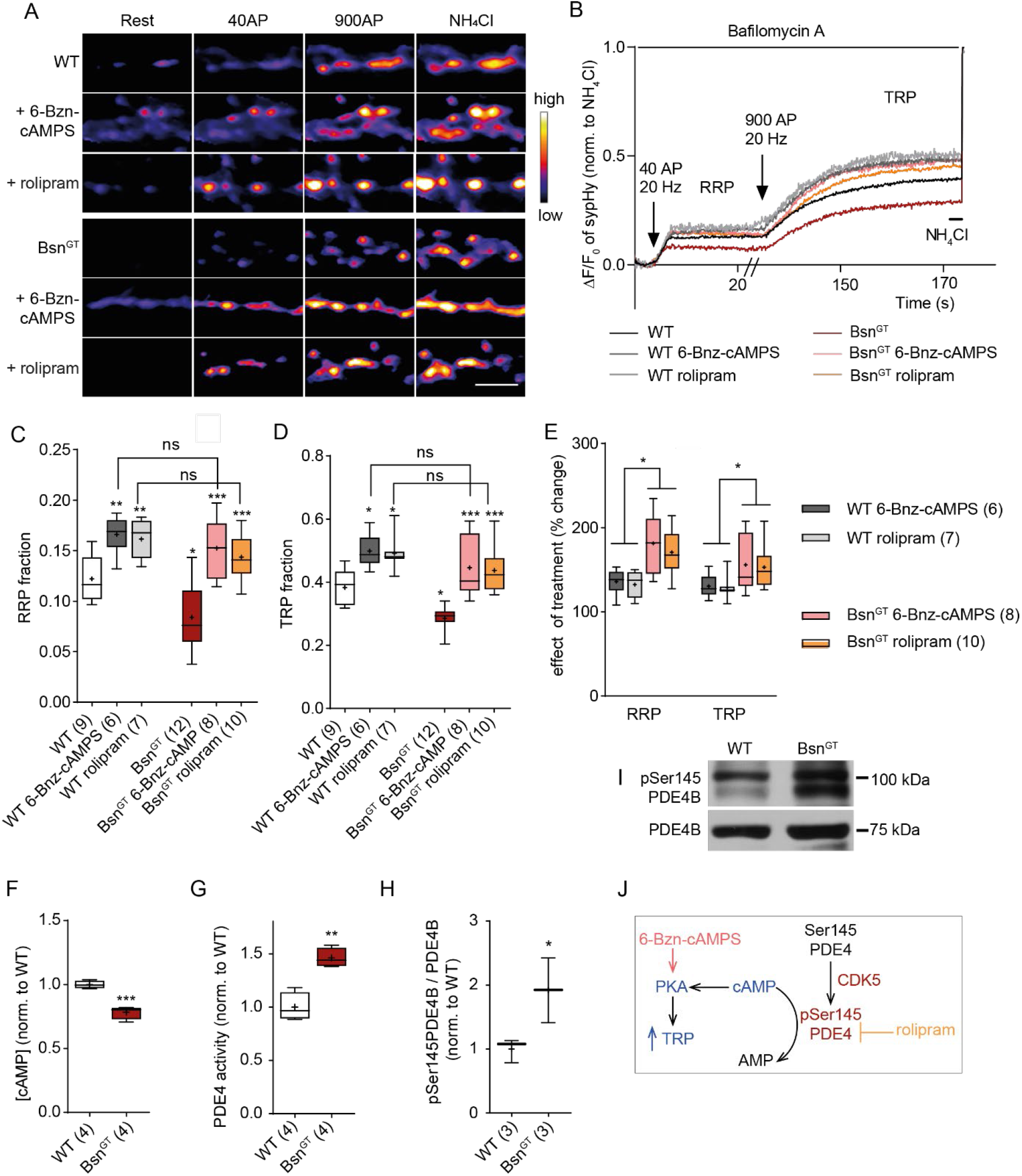
PDE4 activity inhibition rescues Bsn^GT^ phenotype. Representative pseudo colour images (A) and average traces (B) of sypHy fluorescence plotted for untreated WT and Bsn^GT^ neurons (black and red) and cultures treated with of 6-Bnz-cAMPS (50 μM, 1 hour; gray and pink) and rolipram (1 μM, 30 min, light grey and orange). (C, D) Quantifications of RRP and TRP fractions in experiment in A. (E) Effect of treatment is significantly higher in Bsn^GT^ than in WT. (F, G) Quantification of cAMP levels and PDE4-associated phosphodiesterase activity assessed in hippocampal tissue of WT and Bsn^GT^ animals. (I) Representative immunoblot with antibody against CDK5-dependent pSer145PDE4B and total PDE4B on hippocampal tissue from WT and Bsn^GT^ animals. (H) Quantification of pSer145PDE4B to the total PDE4B ratio from experiment in I. (J) The scheme illustrates the CDK5-dependent regulation of cAMP hydrolysis by PDE4 and the downstream effect of PKA affecting the TRP size. The changes in this signalling detected in Bsn^GT^ are shown; blue indicates decreased and red an increased activity or abundance. In the plots, the interquartile range and median are depicted as boxes, minimal and maximal values as whiskers and + indicates mean. The sample size is given in brackets and corresponds to the number of imaging experiments performed on three independently prepared cultures in (C-E) or individual animals used for sample preparation in F-H. Statistical significance was assessed using one-way ANOVA with Tukey’s post hoc test in C and D and Student’s t-test in E, F and H and is given as: *p≤0.05, **p < 0.01, ***p< 0.001. Scale bar is 2μm.

cAMP signalling is terminated by hydrolysis of cAMP by enzymes phosphodiesterases, with PDE4 isoform being the most abundant enzyme isoform in brain. To address a possible role of PDE4 in the regulation SV recycling, we applied the PDE4 inhibitor rolipram (1 μM, 0.5 h) to neurons of both genotypes. This treatment led to an increase in RRP by 39 % and TRP by 29% in WT but had again much stronger effect in Bsn^GT^, where it increased the RRP by 78% and TRP by 56% (Fig 6E). Importantly, similar to Sp-6-Bnz-cAMPS, rolipram treatment fully masked the differences in SV pools between the genotypes (Fig 6A-E; RRP: WT + rolipram 0.16 ± 0.01, Bsn^GT^ + rolipram 0.14 ± 0.01, TRP: WT + rolipram 0.49 ± 0.02, Bsn^GT +^ rolipram 0.44 ± 0.02). These results indicate that increased PDE4-dependent hydrolysis of cAMP underlies the differences in SV recycling in the absence of Bsn. In support of this conclusion, we detected significantly higher PDE4-associated phosphodiesterase activity in the hippocampal tissue of Bsn^GT^ mice compared to their WT littermates (Fig 6G; WT 1.00 ± 0.07, Bsn^GT^ 1.46 ± 0.05).

PKA-dependent phosphorylation of the upstream conserved region 1 (UCR1) of PDE4 is a well-established regulatory mechanism that increases the activity of long PDE4 isoforms (MacKenzie et al., 2002). Therefore, it was surprising to detect high PDE4 activity in the condition of low PKA phosphorylation in Bsn^GT^. Recently, a new CDK5-dependent regulation of PDE4 activity was discovered in neurons. Specifically, CDK5 phosphorylation at Ser145 within the UCR1 (pSer145PDE4B) significantly increased PDE4 phosphodiesterase activity also in the absence of PKA activation (Plattner et al., 2015). Our data indicated an increased presynaptic CDK5 activity in Bsn^GT^ (Fig 3A-E). To test the possible CDK5-dependent activation of PDE4 in Bsn^GT^ we quantified phosphorylation of the CDK5-dependent phosphorylation site in PDE4 (pSer145 PDE4B) in WT and Bsn^GT^ hippocampal homogenates using a phoshphospecific antibody (Plattner et al., 2015). We detected significantly higher levels of CDK5-dependent PDE4B phosphorylation in Bsn^GT^, indicating that activation of PDE4 by its CDK5-dependent phosphorylation drives the exacerbated cAMP hydrolysis observed upon deletion of Bsn (Fig 6H, I; WT 1.00 ± 0.11, n=3 vs Bsn^GT^ 1.92 ± 0.29). Together, these data confirm that reduction in the fraction of releasable SVs in the absence of Bassoon relies on depletion of cAMP due to dysregulation of its PDE4-dependent hydrolysis downstream of CDK5 (Fig 6J). Moreover, they also reveal a new role of CDK5-PDE4-cAMP-PKA signalling axis in the regulation of presynaptic release competence.

## Discussion

### Imaging of SV pools in Bsn^GT^ reveal a decrease in SV release competence

Despite the well-documented impact of Bsn deletion on synaptic neurotransmission, the molecular understanding of this impairment remained limited. In this work, we explored the effect of genetic ablation of Bsn expression on the recycling and release competence of SVs and dissected the cellular signalling involved in this process. We observed a higher fraction of silent excitatory presynapses in Bsn^GT^ cultured neurons compared to controls, which is reminiscent of data obtained by early analyses of mutant mice lacking the central region of the Bsn protein (Altrock et al., 2003). This previous study reported functional inactivation of a fraction of synapses leaving the remaining synapses unaffected and postulated that a subset of synapses relies on normal Bsn expression (Altrock et al., 2003). Our current data modify this statement revealing Bsn as a universally acting factor necessary for proper release competence in glutamatergic synapses. We observed reduced sizes of RRP and TRP of SVs evoked by electrical and chemical stimulation. Reduced RRP has been detected by electrophysiological measurements in synapses of cultured hippocampal neurons, endbulb of Held and ribbon synapses of inner hear cells (IHC), and cone photoreceptors in Bsn mutants (Altrock et al., 2003; Buran et al., 2010; Frank et al., 2010; Jing et al., 2013; Mendoza Schulz et al., 2014; Babai et al., 2019; Babai et al., 2020). In synapses with high-frequency release, such as cerebellar mossy fiber to granular cell synapse, IHC ribbon synapse, and cone photoreceptors, the deletion of Bsn led to more pronounced depression of neurotransmission upon repetitive stimulations (Frank et al., 2010; Hallermann et al., 2010; Jing et al., 2013; Babai et al., 2019; Babai et al., 2020). The stronger depression of neurotransmission was attributed to a slower replenishment of SVs, which is well compatible with a smaller total recycling pool of SVs shown in this study. Thus, the data obtained using the imaging approach in cultured neurons in this study are in good accordance with the previously published data generated by alternative methodologies and using ex vivo preparations.

### Aberrant SV recycling relies on defects in presynaptic phosphor-homeostasis upon Bsn loss

This study revealed a crucial role of Bsn in the maintenance of normal dynamic phosphorylation at presynapse. We measured lower PKA catalytic activity and detected lower phosphorylation of known presynaptic substrates of PKA in the absence of Bsn in vivo and in vitro. The decreased PKA activity was a consequence of dysregulation in the metabolism of cAMP due to its increased hydrolysis and inefficient production. cAMP and its downstream target PKA are the major regulators of presynaptic plasticity (Huang et al., 1994). PKA activation has been linked to the increase in vesicular release probability, but also to an increment in SV release competence and RRP size, which were reduced in Bsn^GT^ mice as shown in previous studies and in this work (Chen and Regehr, 1997; Lonart et al., 1998; Menegon et al., 2006; Moulder et al., 2008; Vaden et al., 2019). cAMP signalling plays a major role in the regulation of SVs at the glutamatergic synapse by neuromodulators (Patzke et al., 2019; Patzke et al., 2021). Therefore, it will be interesting to investigate the effect of Bsn deletion on neuromodulation in future studies. Our experiments also revealed enhanced CDK5-dependent phosphorylation of synaptic targets in Bsn^GT^ neurons. CDK5 regulates the release competence of SVs and drives their assignment to the ResP (Kim and Ryan, 2010, 2013; Verstegen et al., 2014). In line with the increased CDK5 activity, we observed increased ResP in the absence of Bsn. Acute pharmacological activation of PKA or inhibition of CDK5 fully normalized the defect in SV recycling in the absence of Bsn, confirming the pivotal role of the aberrant activity of these enzymes in the decline of SV release competence in Bsn^GT^ neurons. The acute rescue of Bsn^GT^ phenotype also confirms that aberrant presynaptic signalling and not structural or developmental impairments cause the defects in SV recycling in the absence of Bsn. Finally, our experiments revealed a dual role of phosphatase calcineurin in the regulation of SV recycling. Calcineurin antagonizes the effect of CDK5 on the recruitment of SVs into the ResP (Kim and Ryan, 2010, 2013). Compliant with this, calcineurin inhibition decreased the TRP in WT neurons and fully phenocopied the Bsn^GT^ situation indicating that acute shift of CDK5/calcineurin balance in favour of CDK5 induced a fast decrease in release competence of SVs and their assignment to the ResP. Unexpectedly, calcineurin inhibition in Bsn^GT^ but not in WT increased RRP. Calcineurin antagonizes PKA-dependent phosphorylation of Thr138 of SNAP25, which is required for maintenance of RRP in chromaffin cells (Nagy et al., 2004). The selective effect of calcineurin inhibition on RRP in Bsn^GT^ is therefore consistent with elevated calcineurin and with decreased PKA activity that we observed in the absence of Bsn and indicates a role of calcineurin by antagonising PKA in the regulation of RRP in glutamatergic synapses (Fig. 4G).

### PDE4 regulates SV pools

Unexpectedly, an acute pharmacological inhibition of CDK5, which is known to promote the recruitment of SVs to ResP, also affected size of RRP in both WT and Bsn^GT^ neurons. We sought a possible explanation and realised that CDK5 was linked to the regulation of cellular cAMP signalling in the forebrain (Guan et al., 2011; Plattner et al., 2015). Specifically, the phosphorylation by CDK5 was demonstrated to stimulate the phosphodiesterase activity of PDE4, decreasing the cAMP levels, which attenuates the kinase activity of PKA (Plattner et al., 2015). In Aplysia, it was shown that presynaptic PDE4 controls the PKA activity and synaptic facilitation (Park et al., 2005). However, the role of PDE4-dependent hydrolysis of cAMP in the regulation of neurotransmitter release in vertebrate neurons remained elusive. We detected increased CDK5-dependent phosphorylation of PDE4 and enhanced hydrolytic activity of PDE4 in Bsn^GT^. Inhibition of PDE4 activity by rolipram or application of cAMP analogue resistant to PDE4-dependent hydrolysis fully normalized the SV pools in Bsn^GT^ neurons. Thus, we demonstrated that dysregulation of SV pools in the absence of Bsn results from profound dysregulation of presynaptic cAMP signalling. Our data also revealed a new role for PDE4 in the crosstalk of CDK5/calcineurin and cAMP/PKA-dependent regulation of SV recycling and reveal a dual role of presynaptic CDK5 activity that directly reduces SV availability for release by their shift to ResP and indirectly by decreasing RRP via PDE4-mediated inhibition of cAMP signalling. PDE4 has been linked to the pathophysiology of schizophrenia. The product of a schizophrenia risk gene, DISC1, binds and regulates PDE4 activity, which is dysregulated in psychotic patients and in animal models (Millar et al., 2005). Application of selective PDE4 inhibitor rolipram rescued impaired LTP and behavioural deficits in pharmacologically and genetically-induced animal model of psychosis by presynaptic mechanisms indicating a role of PDE4 in presynaptic plasticity (Wiescholleck and Manahan-Vaughan, 2012; Kim et al., 2021). Our work now directly demonstrates an effect of PDE4 on the regulation of SV release competence and thus provides a cellular substrate for the PDE4-dependent modulation of presynaptic plasticity.

### How can Bsn connect to the regulation of dynamic phosphorylation at presynapse?

We have shown previously that Bsn organises presynaptic voltage-gated calcium channels in neuronal cell types (Frank et al., 2010; Hallermann et al., 2010; Ryl et al., 2021). In cortical and hippocampal synapses, it is required for specific recruitment of Cav2.1 (P/Q-type channels) channels to the release sites and consequently, relatively more Cav2.2 (N-type) channels are present at synapses in the absence of Bsn (Davydova et al., 2014). Interestingly, previous studies linked the Cav2.2 to the activity-dependent regulation of presynaptic CDK5/calcineurin activities (Su et al., 2012; Kim and Ryan, 2013). Future studies will be necessary to address whether the dysregulation of presynaptic phosphorylation in Bsn^GT^ originates from aberrant presynaptic calcium influx in the absence of Bsn. Another possible explanation of the effect of Bsn deletion on presynaptic phosphorylation status described in this work is a potential role of Bsn in scaffolding of presynaptic kinase/phosphatase signalling complexes. The kinase scaffolding proteins such as 14-3-3s or AKAPs emerged to increase selectivity and efficiency of substrate recognition and recruitment (Fu et al., 2000; Torres-Quesada et al., 2017). Indeed, Bsn interacts with 14-3-3s, which can recruit signalling complexes to the active zone (Schroder et al., 2013). Moreover, Bsn emerged as a heavily phosphorylated protein and several studies revealed significant changes in its phosphorylation pattern during neuronal activity (Kohansal-Nodehi et al., 2016; Engholm-Keller et al., 2019; Silbern et al., 2021). Thus, Bsn might act as a docking site for multiple signalling complexes at presynapse and organise the neuronal-activity dependent presynaptic signalling. Future work will be necessary to explore the individual molecular interactions behind the Bsn-linked regulation of presynaptic kinase/phosphatase dynamics.

## Materials and Methods

### Animals

Bassoon knockout mice (Bsn^GT^) were generated from Omnibank ES cell line OST486029 by Lexicon Pharmaceuticals, Inc. (The Woodlands, TX) as reported previously (Hallermann et al., 2010). In this strain, a functional inactivation of Bsn allele relies on gene trap strategy. To get the respective Bsn^GT^ and WT animals for our studies, heterozygous animals were backcrossed for 8-10 generations to C57BL/6N background and then bred at a 2:1 ratio. For primary neuronal culture preparations and all other experiments, animals of both sexes were used. Breeding of animals and experiments using animal material was carried out according to the European Communities Council Directive (2010/63/EU) and approved by the local animal care committees of Sachsen-Anhalt, Germany.

### Antibodies

The primary antibodies were used for immunocytochemistry (ICC), Western Blot (WB), and Syt1Ab uptake in the concentration indicated as follows: rabbit antibodies against Syt1 (labelled with Oyster 550; live staining: 1:70, # 105103C3, Synaptic System, Göttingen, Germany), pThr138SNAP25 (WB:1:500, # 042077, Biomol), SNAP-25 (WB: 1:1000, # 111002, Synaptic System), pSer9Syn1 (ICC:1:1000, gift from Dr. Fabio Benfenati, IIT, Genova, Italy), pSer551Syn1 (ICC: 1:1000, WB:1:1000, gift from Dr. Anna Fassio, IIT, Genova, Italy) pSer9Syn1 (WB: 1:1000, # 2311S, Cell Signaling), PKAα cat (C-20) (WB:1:500, # sc-903, Santa Cruz), pSer145PDE4B (WB 1:200, (Plattner et al., 2015)), mouse antibodies against Syn1 (ICC: 1:1000, WB: 1:1000, # 106011, Synaptic System), VGLUT1 (ICC:1:1000, # MAB5502, Millipore), guinea pig antibodies against Syn1 (ICC:1:1000, # 106104, Synaptic System), VGAT (ICC:1:1000, # 131004, Synaptic System) and sheep antibody against PDE4B (WB 1:500, (Huston et al., 1997)). Secondary antibodies used for ICC were: donkey anti mouse coupled with Alexa 488-(ICC 1:2000, # A21202, Thermo Fisher Scientific), Cy3-donkey anti rabbit (1:2000, #711-165-152, Dianova/Jackson ImmunoResearch Labs), and Cy5-donkey anti guinea pig (1:1000, # 706-175-148, Dianova/Jackson ImmunoResearch Labs). For fluorescent detection of WB, IRDye 680 donkey anti mouse # 926-68072 and IRDye 680RD donkey anti rabbit # 926-68071 and secondary antibodies from Li-COR were used. For chemiluminescent detection, Thermo Fisher #31460 anti-rabbit; #A16047 anti-sheep HRP conjugated antibodies were used.

### Chemical reagents

D-(−)-2-amino-5-phosphonopentanoic acid (APV) (CAS #79055-68-8), 6-cyano-7-nitroquinoxaline-2,3-dione disodium (CNQX) (CAS #479347-85-8), KN-93(CAS #139298-40-1), (R)-(−)-Rolipram (CAS # 85416-75-7), TTX (CAS #4368-28-9), and (+)-Bicuculline (CAS #485-49-4) were purchased from Tocris. H-89 (CAS #130964-39-5). Forskolin (CAS #66575-29-9) and FK-506 monohydrat (CAS #109581-93-3), were from Sigma-Aldrich. InSolution Roscovitine (CAS # 186692-46-6) and Bafilomycin A1 (CAS #88899-55-2) were from Calbiochem. N6-Benzoyladenosine-3’, 5’-cyclic monophosphorothioate, Sp- isomer (Sp-6-Bnz-cAMPS), sodium salt was from Biolog (CAS # 152218-18-3)

### Cultures of primary hippocampal neurons

Primary hippocampal cultures from neonatal (P0-P1) Bsn^GT^ and their WT littermates were prepared as described previously (Altmuller et al., 2017; Montenegro-Venegas et al., 2021). Briefly, hippocampal tissue was treated with 0.25% (final c) of trypsin (# 15090046) and cell suspension was obtained by mechanical trituration. Cells were plated in densities of 35000 cells per coverslip (18 mm diameter) on poly-L-lysine coated coverslips. One hour after plating, coverslips were transferred with neurons facing down into dishes containing 60-70% confluent monolayer of astrocytes and Neurobasal A medium (#12349015) supplemented with 2% (v/v) B27 (#17504044), 1 mM sodium pyruvate (# 11360070), 4 mM GlutaMAX™ supplement (# 35050038) and antibiotics (100 U/ml penicillin, 100 μg/ml streptomycin, # 15240096). About 1 mm large paraffin dots were placed onto coverslips prior to coating and cell plating to assure medium exchange between feeder layer and neurons during all time of culturing. All chemicals used for neuronal cultures were obtained from Thermo Fisher Scientific unless indicated otherwise. To prevent overgrowth of astroglia 0.6 μM AraC (CAS # 147-94-4, Sigma Aldrich) was added to the cells at day 1 and 3 after plating to reach a final concentration of 1.2 μM. All cells were maintained at 37°C in a humidified incubator containing 5% CO_2_.

### Quantitative immunostaining and image analysis

Quantitative immunostainings were performed, acquired and analysed as described previously with minor differences (Lazarevic et al., 2011). For staining, all samples compared in one experiment were processed in parallel with identical solutions. Image acquisition was done for all samples from one experiment on the same day with identical camera and illumination settings. For IF quantification, 7-10 cells from 2 independent coverslips were acquired and quantified for each experiment in odder to reduce experimental variance. Unspecific background signal was removed using threshold subtraction in ImageJ software (NIH, http://rsb.info.nih.gov/ij/). In all experiments, synaptic puncta were defined semiautomatically by setting rectangular regions of interest (ROI) with dimensions of about 0.8 × 0.8 μm around local intensity maxima in the image with staining for synaptic marker Syn1, VGLUT1, or VGAT using OpenView software written and kindly provided by N.E. Ziv (Tsuriel et al., 2006). Mean IF intensities were measured in synaptic ROIs in all corresponding channels using the same software and normalized to the mean IF intensities of the control group in each experiment. For analyses of active synapses, puncta with over-threshold staining for Syn1 and VGLUT1 or VGAT were considered as excitatory or inhibitory synapses.

### Synaptotagmin1 antibody uptake

For quantitative assessment, all coverslips compared in one experiment were processed in parallel using identical antibodies solutions and other reagents as described earlier (Lazarevic et al., 2011). For the Syt1Ab uptake, neurons grown for 19-21 days in vitro (DIV) were washed with Tyrode’s buffer (TB) containing:119 mM NaCl, 2.5 mM KCl, 2 mM CaCl_2_, 2 mM MgCl_2_, 30 mM glucose, and 25 mM HEPES, pH 7.4, and then incubated with Oyster 550-labeled antibody against luminal domain of Syt1. To monitor endogenous network activity-induced Syt1Ab uptake neurons were incubated with Oyster 550-labeled anti Syt1 antibody in TB for 20 minutes at 37°C. To visualise all release-competent vesicles Oyster 550-labeled anti Syt1 antibody was applied to cells in high K^+^ TB containing 71.5 mM NaCl, 50 mM KCl, 2 mM CaCl_2_, 2 mM MgCl_2_, 30 mM glucose, and 25 mM HEPES, pH 7.4 for 4 min. Then cells were washed twice with TB, fixed with 4% (w/v) PFA, permeabilized with 0,3% (v/v) Triton X-100, and stained with specific antibodies as described earlier (Lazarevic et al., 2011). Synapses with over-threshold IF for Syt1Ab uptake were considered as active synapses.

### Lentiviral particles production

The lentiviral vector for expression of ratio:sypHy (Rose et al., 2013) was described in (Lazarevic et al., 2017). Lentiviral particles were generated in HEK293T cells (ATTC, Manassas, VA, USA) using psPAX2 packaging and pVSVG pseudotyping vectors (Lois et al., 2002). HEK293T cells were grown in media containing 10 % (v/v) FCS to 60 % confluence in the 75 cm2 flasks. Cells were transfected with 20 μg of total DNA per flask using calcium phosphate method (Fejtova et al., 2009). Molar ratio of FUGW: psPAX2: pVSVG was 4:2:1. 6-8 h after transfection, medium was changed to 10 ml production medium containing Neurobasal A supplemented with antibiotics, 1 mM sodium pyruvate (Life Technologies), B27 and 1mM GlutaMAX™ (Life Technologies). 48 h after transfection, virus-containing media was collected and cleared off large cellular debris by centrifugation for 20 min at 2000 g. Virus-containing supernatant was aliquoted and stored at −80°C.

### Synapto-pHluorin (sypHy) Imaging

Imaging experiments for assessment of RRP and TRP were carried out in 19-20 DIV hippocampal neurons. Cells were transduced with the lentiviral particles expressing ratio:sypHy at day 3-4. Transduced neurons were identified by their RFP expression. Coverslips were mounted in a chamber equipped with field stimulation electrodes (RC-49MFSH; Warner instruments). Electrical stimulation was generated using an isolator unit (WPI) controlled by Master-8 pulse generator (AMPI). During whole experiment cells were kept in TB supplemented with 50 μM APV and 25 μM CNQX to avoid the recurrent network activity-driven release and with 1 μM of Bafilomycin A to prevent the vesicle re-acidification of once released vesicles (Sankaranarayanan and Ryan, 2001). Imaging experiments were done at RT on an inverted microscope (Zeiss Axio Observer.A1) using 63x oil immersion objective (NA 1.4) and GFP/mCherry ET filter set (single-band exciters 470/40 and 572/35, dichroic Chroma #59022BS, emitter Chroma #59022m). Images were acquired at frame rate of 12 Hz using an EMCCD camera (Evolve 512; Photometrics) controlled by VisiView (Visitron Systems GmbH). After 5 s of baseline acquisition (F_0_), neurons were stimulated with 40 AP at 20 Hz to release of RRP. After 2 min recovery time, TRP was released by application of 900 AP at 20 Hz, 1 min later a pulse of TB containing 60 mM NH_4_Cl was applied to visualise all sypHy expressing vesicles (Burrone et al., 2006).

For analysis, first background (calculated form 4 independent background regions within the image) was mathematically subtracted. Responding synaptic puncta were visualised by subtracting the first 10 frames corresponding the baseline from the traces obtained directly after the 900 AP stimulus. Circular ROIs with a diameter of 8 × 8 pixels were placed over each responding bouton and mean IF values from between 80 and 150 ROIs per visual field were measured using the Time Series Analyzer V2.0 plugin in ImageJ (NIH, http://rsb.info.nih.gov/ij/). Only the boutons showing stable responses and TRP ≤80% of NH_4_Cl-evoked fluorescence were considered for the analyses. Bleaching correction was done by normalizing to the bleaching factor obtained from imaging experiments without stimulation performed on the same coverslip. Traces were plotted using GraphPad software. The relative sizes of the RRP and the TRP were expressed as fractions of the total sypHy-expressing pool detected after the addition of NH_4_Cl (F_NH4Cl_). RRP and TRP were quantified by averaging the mean of 100 values per ROI plateau phases (frames150–250 corresponding to time points between 10 and 20 sec after start of image acquisition and 600-700 corresponding to time points ≈165-175 sec, respectively, on the XY graph). ResP was quantified at as F_NH4Cl_ – F_TRP_.

### Brain fractionation and quantitative Western blot

Hippocampi were isolated from 8 weeks old WT and Bsn^GT^ littermates of both sexes. Fresh tissue was homogenized in buffer containing 25 mM Tris-HCl pH 7.4, 0.32 M sucrose supplemented with Complete mini protease inhibitors (# 11836153001, Roche), PhosStop phosphatase inhibitor cocktail (# 4906837001, Roche) using Potter glass-Teflon homogeniser (B. Braun International) with 12 strokes at 900rpm. Cell debris and nuclei were sedimented at 1000xg for 10 min at 4°C. The supernatant was collected and centrifuged again at 12000xg for 20 min at 4°C to obtain supernatant (S2) corresponding to cytoplasmic fraction and pellet (P2) corresponding to the crude membrane fraction. Protein concentrations in fractions were determined using the BC assay (# UP40840A, Interchim).

To obtain samples for Western blotting, the P2 pellet was resuspended in 25 mM Tris-HCl, pH 7.4, 150 mM NaCl, 1% (v/v) Triton X-100 supplemented with Complete mini protease inhibitors and PhosStop phosphatase inhibitor cocktail. An equal amount of proteins (20 μg/lane) were separated using one-dimensional SDS-PAGE and then transferred to Millipore Immobilon-FL PVDF membranes. For immunodetection the primary antibody were applied at 4°C overnight and the fluorescently-labelled secondary antibodies for 1 h at room temperature. Antibodies were diluted in PBS containing 0.1% (v/v) Tween 20, 5% (w/v) BSA, and 0.025% (w/v) sodium azide. Immunodetection was carried out using chemiluminescent imager for quantification of pSer145PDE4B and PDE4B immunoreactivity and using Odyssey Infrared Imagine System and Odyssey software v2.1 (Li-COR) for all other experiment. Quantification of the blots was performed using ImageJ software. After background subtraction, all values were normalized to respective loading controls.

### Measurement of PKA activity

P2 pellets were prepared from hippocampi of 8 weeks old animals as described above and resuspended in a buffer containing 25mM Tris-HCl pH 7.4,150 mM NaCl, 0.5% (v/v) Triton-X100 and Complete mini protease and PhosStop phosphatase inhibitor cocktail. Samples containing equal protein amount, as assessed by BC assay (#UP40840A, Interchim), were incubated with GammaBind Plus Sepharose beads (#GE17-0886-02, GE Healthcare) coupled with rabbit polyclonal antibody against PKAα cat (C-20; Santa Cruz Biotechnology) for 4 h at 4°C. After washing three times with PKA assay buffer (40 mM Tris pH 7.5, 20 mM MgCl_2_, 0.1 mg BSA, 0.1 mM DTT) the immunoprecipitates were resuspended in 50 μl of the same buffer supplemented with 70 μM ATP and 0.3 mM Kemptide, a PKA substrate (LRRASLG; # V5601 Promega) and incubated for 0.5 h at 30°C in 96-well plate. The ATP hydrolysis was detected upon addition of 50 μl/well luminescent Kinase Glo plus reagent (# V6711, Promega) using FLUOstar Omega microplate reader (BMG Labtech).

### Calcineurin activity assay

Calcineurin activity was assessed using Calcineurin activity assay kit (#20700, Calbiochem). Hippocampus tissue of 8 weeks old animals was homogenised in the lysis buffer (25 mM Tris-HCl pH 7.5, 50 μM EDTA, 50 μM EGTA, 0.2% (v/v) Nonidet P-40, 0.5 mM dithiothreitol, Complete mini protease inhibitors) by gently pressing through 16G needle. Homogenates were cleared by centrifugation at 100.000g at 4°C for 45 min. Protein concentration was assessed by BC assay and samples with an equal protein amount were processed for the assay. Free phosphate was removed, samples were rebuffered to the assay buffer containing 100 mM NaCl, 50mM Tris-HCl (pH 7.5), 6 mM MgCl_2_, 0. 5mM CaCl_2_, 0.5 mM dithiothreitol, 0.05% (v/v) Nonidet P-40 and incubated with RII phosphopeptide DLDVPIPGRFDRRVpSVAAE, a specific calcineurin substrate, for 0.5 h at 30°C in a 96-well plate. The phosphate release was detected upon addition of 100 μl/well of Green reagent by absorbance measurement at 620 nm. The Ca^2+^-independent phosphatase activity was measured in samples supplemented with EGTA and subtracted from measured values to obtain calcineurin phosphatase activity.

### cAMP assay

P2 pellets were prepared from hippocampi of 8 weeks old animals as described above and dissolved in buffer containing 0.1M HCl and 0.5% Triton-X 100. The cAMP levels in samples containing equal protein concentrations were quantified using ELISA-based assay kit (#ADI-900-163, ENZO Life Sciences). The assay was done following the manufacturer’s instructions.

### PDE4 immunoprecipitation and ictivity assay

PDE4 activity was assessed using a cyclic nucleotide phosphodiesterase assay kit (#BML-AK800-0001, Enzo Life Sciences) according to the supplier’s instructions. In brief, hippocampal crude membrane pellets P2 prepared as described above, were resuspended with ice-cold buffer contained 50 mM HEPES pH 7.4, 120 mM NaCl, 0.1 mM EDTA, 0.5% (v/v) Triton X-100 and Complete mini protease inhibitors (Roche). Upon removal of free phosphate, samples containing same protein amount were subjected to immunoprecipitation with rabbit polyclonal antibody against PDE4B (H-56) (#sc-25812, Santa Cruz Biotechnology) coupled to GammaBind Plus Sepharose beads (# 17-0886-02, GE Healthcare) for 4 h at 4°C. After three times washing with PDE assay buffer (10 mM Tris-HCl, pH 7.4), immunoprecipitates were resuspended in 50 μl of the same buffer supplemented with 0.2 mM 3’,5’-cAMP substrate and 5’-nucleotidase enzyme (50 kU/well) and incubated for 0.5 h at 30°C in a 96-well plate. Phosphate release was detected upon incubation with Green reagent (100 μl/per well) for 30 min by measurement of absorbance at 620 nm in a microplate-reading spectrophotometer.

### Quantification and statistical analysis

All results of quantitative analyses are given as means ± standard errors of the mean (SEM). Statistical analyses were performed with Prism 8 software (GraphPad Software, Inc.) using Student’s t-test or one-way ANOVA as indicated for each experiment. Statistical significance is marked as *P < 0.05, **P < 0.01, ***P < 0.001 in all plots. Information about group character and size is specified in the figure legend for each experiment.

## Acknowledgment

We thank Janina Juhle, Juliana Monti, Sabine Müller, Peggy Patella, and the animal facility at LIN Magdeburg and PETZ Erlangen for excellent technical assistance. We also acknowledge Anna Fassio, Fabio Benfenati, James A. Bibb, and Miles D. Housley for generously providing the non-commercial antibodies used in this study.

## Funding

This study was supported by DFG (FE1335/3) and BMFB GeNeRARe (FZ 01GM1902B) to AF, ERANET/BMBF (AMRePACELL) and BMBF 01DN17002 (Preplastic) to EDG, Center for Behavioural Brain Sciences—CBBS promoted by Europäische Fonds für regionale Entwicklung—EFRE (ZS/2016/04/78113) to CMV.

## Conflicts of interest/Competing interests

The authors declare that they have no conflict of interest

## Ethics approval for experiments using animal material

Animal experimentation: Breeding of animals and experiments using animal material were carried out in accordance with the European Communities Council Directive (2010/63/EU) and approved by the local animal care committees of Sachsen-Anhalt.

## Authors’ contributions

Conceptualization, supervision: CMV, AF; formal analysis, project administration, data curation, writing - original draft: CMV, AF; investigation: CMV, DG, FP; methodology: CMV, EPF, MAA, FP, VL, AF; funding acquisition: CMV, EDG, AF; resources: EDG, AF; visualization: CMV, DG, AF; writing - review & editing: all authors

## Notes

### Competing Interest Statement

The authors have declared no competing interest.

